# Antisocial rewarding in structured populations

**DOI:** 10.1101/092932

**Authors:** Miguel dos Santos, Jorge Peña

## Abstract

Cooperation in collective action dilemmas usually breaks down in the absence of additional incentive mechanisms. This tragedy can be escaped if cooperators have the possibility to invest in reward funds that are shared exclusively among cooperators(prosocial rewarding). Yet, the presence of defectors who do not contribute to the public good but do reward themselves(antisocial rewarding) deters cooperation in the absence of additional countermeasures. A recent simulation study suggests that spatial structure is sufficient to prevent antisocial rewarding from deterring cooperation. Here we reinvestigate this issue assuming mixed strategies and weak selection on a game-theoretic model of social interactions, which we also validate using individual-based simulations. We show that increasing reward funds facilitates the maintenance of prosocial rewarding but prevents its evolution from rare, and that spatial structure can sometimes select against the evolution of prosocial rewarding. Our results suggest that, even in spatially structured populations, additional mechanisms are required to prevent antisocial rewarding from deterring cooperation in public goods dilemmas.

## Introduction

Explaining the evolution of cooperation has been a long-standing challenge in evolutionary biology and the social sciences^1–6^. The problem is to explain how cooperators, whose contributions to the common good benefit everybody in a group, can prevent defectors from outcompeting them, leading to a tragedy of the commons where nobody contributes and no common good is created or maintained^7^.

A solution to this problem is to provide individuals with additional incentives to contribute, thus making defection less profitable^8,9^. Incentives can be either negative (punishment) or positive (rewards). Punishment occurs when individuals are willing to spend resources in order for defectors to lose even more resources^10^. Punishment can be stable against defection, since rare defectors are effectively punished^11^. However, to evolve from rare and resist invasion by individuals who cooperate but restrain from investing into incentives, i.e., second-order defectors, punishers must gain from punishing, for example, through reputational benefits in future interactions^9,12,13^.

Defection in collective action problems can also be prevented via positive incentives. Using rewards, cooperators can pay to increase the payoff of other cooperators. While the emergence of such behavior is usually favored, as there are very few cooperators to reward when cooperators are rare, it becomes increasingly costly to sustain as cooperators become more abundant in the population^14^. To resist second-order defectors, non-rewarding players must benefit less from rewards^15^. Alternatively, when both rewards and punishment are present, rewards can foster the emergence of punishment, which in turn can be stable provided second-order punishment is available^16^.

Individuals can either decide to impose incentives unilaterally, or they can pool their effort to impose incentives collectively. When acting collectively, individuals can be thought as investing into a fund used to either punish defectors or reward cooperators; in the latter scenario one speaks of “prosocial rewarding”. These collective mechanisms can be viewed as primitive institutions, as group members both design and enforce the rules to administer incentives to overcome social dilemmas^17^. Pool rewards^15,18^ are particularly interesting because they involve the creation of resources, as opposed to their destruction (as in punishment). Prosocial rewarding can favor cooperation only if non-rewarding players can be sufficiently prevented from accessing reward funds so that second-order defectors benefit less from rewards than do rewarders^15^. However, the presence of “antisocial rewarders”, i.e., individuals who do not contribute to the public good but reward themselves, destroys cooperation unless additional mechanisms, such as better rewarding abilities for prosocials, work in combination with exclusion^19^.

Pool rewards can also be viewed as a second collective action dilemma played exclusively among those players who made a similar choice in the first public goods game, i.e., cooperators with each other, and defectors with each other. The nature of this secondary collective action is not necessarily similar to that of the first public goods game, and might, for example, involve non-linear returns. This observation extends beyond human behavior, with situations where individuals are involved in different levels of social dilemmas being particularly likely in bacterial communities^20^. Indeed, many species of bacteria secrete public good molecules (e.g., iron-binding siderophores and other signaling molecules), which are susceptible to exploitation from both their own and other strains^21–23^. In addition, bacteria are also involved in within-species public goods games, as some of those public good molecules can also be strain-specific^23,24^. Thus, the relevance of pool-reward mechanisms extends to non-human species.

A recent theoretical study by Szolnoki and Perc^25^ [hereafter, SP15] challenged the view that additional mechanisms are required to prevent antisocial rewarding from deterring cooperation in public goods games. Contrastingly, SP15 showed that,if individuals interact preferentially with neighbors in a spatially structured population, prosocial rewarding outcompetes antisocial rewarding and that increasing rewards is beneficial for prosocial rewarding. However, SP15 reached these conclusions by means of Monte Carlo simulations of a very specific model of spatial structure and evolutionary dynamics (a square lattice with overlapping groups and a Fermi update rule) where interacting groups are always equal to five. Hence, it remains unclear whether their results generalize to a broader range of spatial models.

Spatial structure can favor the evolution of cooperation. The main reason for this phenomenon is simple. When populations are spatially structured through limited dispersal, social interactions necessarily occur more often among relatives. Hence, kin selection is at work^3,26–28^. However, spatial structure also means that competitors are also more often kin^29^. In certain models (such as an island model with Wright-Fisher demography^30^ or an evolutionary graph updated with a Moran birth-death process^31,32^), these two effects cancel each other out and spatial structure has no effect on the evolution of cooperation. More generally, the net effect is not null, and its direction and magnitude can be often conveniently captured by a single “scaled relatedness coefficient”^33^. The scaled relatedness coefficient depends on the demographic assumptions of a given model, including the “update rule” used to implement the evolutionary dynamics, but is otherwise independent of the payoffs from the game used to model social interactions^33–37^.

Studies on spatial games and evolutionary graph theory have also investigated the effects of spatial structure on evolutionary game dynamics^38–41^. Theses studies have shown how particular features of the graph used to represent the population and the update rules can promote or hinder the evolution of cooperation. In particular, it has been shown that, assuming weak selection on discrete strategies and additive effects, the interplay between graph topology and update rule can be captured by a single “structure coefficient” independent of the underlying game^42,43^. Importantly, the structure coefficient of evolutionary graph theory and the scaled relatedness coefficient of kin selection theory are connected by a simple transformation^35^. Therefore, using scaled relatedness as a measure of spatial structure allows one to capture a large variety of spatial models, including spatial games and evolutionary graphs.

Here, we formulate a mathematical model that clarifies the role of spatial structure for cooperation to be favored through pool rewarding. In contrast to SP15 [which, in the tradition of spatial games and evolutionary graphs^38,40,41,44^, assume discrete strategies and strong selection] we assume continuous strategies and weak selection^45,46^. Our different assumptions allow us to build on existing theoretical work^36^ to analytically derive the conditions under which cooperation is favored and to write them as functions of the parameters of the game (including the group size) and of a single “scaled relatedness coefficient”^33,35,36^, which serves as a natural measure of spatial structure. This allows us to make general predictions about the effect of spatial structure on cooperation, and to make connections between our results and the vast literature on social evolution theory^3,28,47^.

## Model

### Public goods game with prosocial and antisocial reward funds

We consider a collective action problem with an incentive mechanism based on reward funds following the model of SP15. Individuals interact in groups of size *n* and play a linear public goods game (PGG) followed by a rewarding stage with non-linear returns. There are two types of actions available to individuals: “rewarding cooperation” (*RC*, or “prosocial rewarding”), whereby a benefit *r*_1_*/n* is provided to all group members (including the focal) at a cost *γ*, and “rewarding defection” (*RD*, or “antisocial rewarding”), whereby no benefit is provided and no cost is payed. The parameter *r*_1_ is the multiplication factor of the PGG, and it is such that 1 < *r*_1_ < *n*; this ensures that the first stage of the game is a prisoner’s dilemma, so that if rewards are absent rewarding cooperation is a dominated strategy.

Individuals choosing *RC* or *RD* also invest in their own reward funds. Each reward fund yields a per capita net reward *r*_2_ −*γ* (reward benefit *r*_2_ minus cost of contributing to the reward pool *γ*) provided there is at least another individual playing the same action among the *n* − 1 other group members, and zero otherwise (i.e., self-rewarding is not allowed and the cost *γ* is payed only if the rewarding institution is created). For example, a focal individual playing *RC* will pay the cost and receive the reward only if there is at least another *RC* among its *n* − 1 partners. This reflects a situation where reward funds yield non-linear returns, and is reminiscent of those of a volunteer’s dilemma^48^. Since the net reward *r*_2_ − *γ* does not depend on the group size *n*, *r*_2_ can in principle take any value greater than, or equal to *γ*. Note that individuals choosing the most common action are more likely to get the reward, even under random group formation. Hence, *RC* can prevail as long as its frequency in the global population is above one half and rewards outweigh the net cost of contributing to the PGG. However, if self-rewarding is allowed, cooperation is never favored even when all individuals play *RC*^19^.

With the previous assumptions, and letting without loss of generality *γ* = 1, the payoffs for a focal individual choosing either *RC* or *RD* when *k* co-players choose *RC* (and *n* −1 −*k* co-players choose *RD*) are respectively given by (cf. Equations 3.1, 3.2, and 3.3 in SP15):

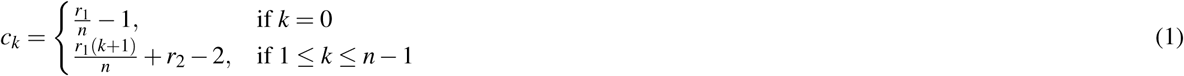

and

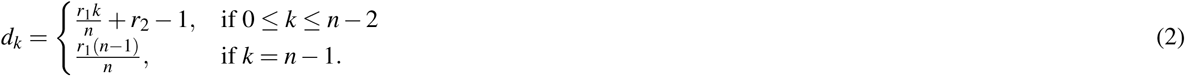

Note that if everybody plays *RC*, everybody gets a payoff *c_n_*_−1_ = *r*_1_ + *r*_2_ − 2. Instead, if everybody plays *RD*, everybody gets *d*_0_ = *r*_2_ −1. Since *r*_2_ *>* 1, *c_n_*_−1_ *> d*_0_ holds for all values of *r*_1_ and *r*_2_, which means that full prosocial rewarding Pareto dominates full antisocial rewarding: Players are collectively better if all play prosocial rewarding with probability one rather than if all play antisocial rewarding with probability one. Therefore, despite the presence of rewards available to both cooperators and defectors, the game we study retains the characteristics of a typical social dilemma where full cooperation by all individuals in the group yields higher payoffs than full defection by all individuals in the group. In such situations, it is usually expected that spatial structure facilitates the evolution of *RC*. As we show below, this is not always the case.

### Spatial structure and evolutionary dynamics

We consider a homogeneous spatially structured population of constant and finite size *N*_T_ where individuals interact with *n* − 1 other individuals according to the game described above. The exact type of spatial structure can follow any of a large family of models, including variants of the island model^49^ and transitive evolutionary graphs^50^. In this last case, and for simplicity, we assume that individuals play a single (*k*+1)-player game with their *k* nearest neighbors, where *k* is the degree of the graph. All that is required for our analysis to be valid is that the selection gradient can be written in a form proportional to the gain function given by Eq. (3) below. We refer the interested reader to previous literature^28,33–36^ for more details on this formalism and the models of spatial structure captured by our approach.

We assume that individuals implement mixed strategies, i.e., they play *RC* with probability *z* and *RD* with probability 1 − *z*, and investigate the evolutionary dynamics of the phenotype *z*. More specifically, we consider the fixation probability *ρ*(*z, δ*) of a single mutant playing *z* + *δ* in a resident population of phenotype *z*, take the phenotypic selection gradient 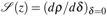 as a measure of evolutionary success^35,51^, and look into convergence stable strategies^52^ under trait substitution dynamics^28^.

In order to evaluate the selection gradient, we make use of standard results regarding the evolution of a continuous phenotype in a spatially structured population^28^. Denoting by *z*_•_ the phenotype of a focal individual, by *z*_o_ the average phenotype of the individuals it socially interacts with, and by *f*(*z*_•_, *z*_o_) the fecundity of the focal individual, and further assuming that fecundity is proportional to the expected payoffs from the game, the selection gradient 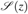 can be shown to take the form^34^

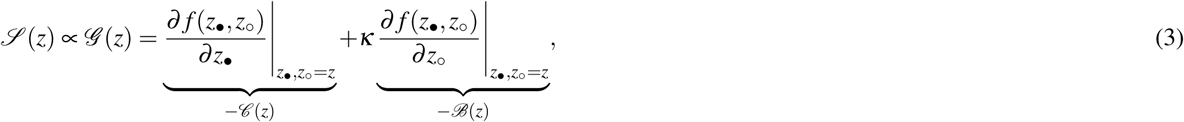

where 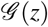 is the “gain function”, which consists of three components: (i) the effect of the focal individual’s behavior on its fecundity (i.e., the “direct effect” 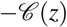, (ii) the effect of the co-players’ behavior on the focal individual’s fecundity (i.e., the “indirect effect” 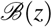, and (iii) a measure of relatedness between the focal individual and its neighbors, demographically scaled so as to capture the effects of local competition (i.e., the “scaled relatedness coefficient” *κ*).

For the matrix game with two pure strategies as the one we consider here, the direct and indirect effects appearing in Eq. (3) can be written, up to a constant factor, as^36^

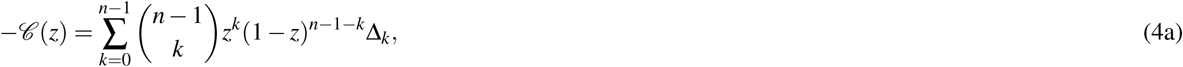

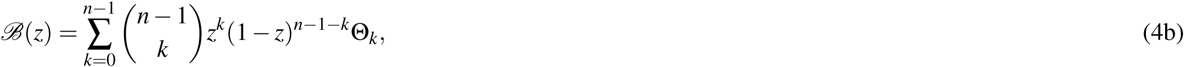

where

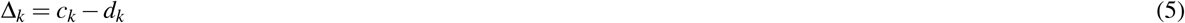

are the “direct gains from switching” recording the changes in payoff experienced by a focal if it unilaterally switches its action from *RD* to *RC* when *k* co-players stick to *RC* and *n* − 1 − *k* stick to *RD*^53^, and

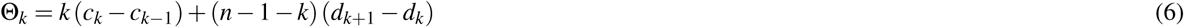

are the “indirect gains from switching” recording the changes in the total payoff accrued by co-players when the focal unilaterally switches its action from *RD* to *RC*^36^. Eq. (4) expresses 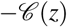 and 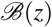 as expected values of the direct and indirect gains from switching when the number of other individuals playing *RC* is distributed according to a binomial distribution with parameters *n* and *z*.

A necessary and sufficient condition for a mutant with phenotype *z* + ***δ*** to have a fixation probability greater than neutral when ***δ*** is vanishingly small is that 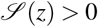 and hence that 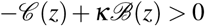 holds. This condition can be interpreted as a scaled form of Hamilton’s rule^33,35^. Importantly, the gain function 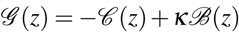 allows one to identify “convergence stable” evolutionary equilibria^54–56^; these are given either by singular strategies *z*\* (i.e., the zeros of the gain function) satisfying 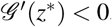, or by the extreme points *z* = 0 (if 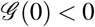) and *z* = 1 (if 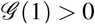). Convergence stability is a standard way of characterizing long-term evolutionary attractors; a phenotype *z*\* is convergence stable if for resident phenotypes close to *z*\* mutants can invade only if mutants are closer to *z*\* than the resident^52^.

### Scaled relatedness

The coefficient *κ* appearing in Eq. (3) is the “scaled relatedness coefficient”, which balances the effects of both increased genetic relatedness and increased local competition characteristic of spatially structured populations^29,33–35,37^. The scaled relatedness coefficient has been calculated for many models of spatial structure for which Eq. (3) applies, see Table 2 of ref.^33^, Table 1 of ref.^34^, Appendix A of ref.^36^, and references therein for some examples. We also note that by identifying

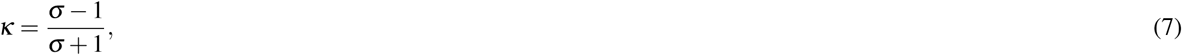

where *σ* is the so-called “structure coefficient”, the right hand side of Eq. (3) recovers the “canonical equation of adaptive dynamics with interaction structure” (ref.^43^, Eq. 5). Table 1 of ref.^43^ provides several examples of models of spatial structure and their respective values of *σ*; transforming these values via Eq. (7), the scaled relatedness coefficients for such models can be obtained in a straightforward manner.

Usually, *κ* takes a value between −1 and 1 depending on the demographic assumptions of the model, but it is always such that the larger it is the less genetic relatedness is effectively reduced by the extent of local competition. Importantly, the larger the magnitude of scaled relatedness *κ* the more important the role of the indirect effect 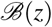 in the selection gradient. For this reason, we use scaled relatedness *κ* as a measure of spatial structure; we hence refer in the following to an increase in *κ* as an increase in spatial structure.

In the following, we present some examples to illustrate the connection between explicit spatial structure models and the formalism we use here, based on the scaled relatedness coefficient. For a well-mixed population or an island model with Wright-Fisher demography (such that generations are overlapping), the value of scaled relatedness is

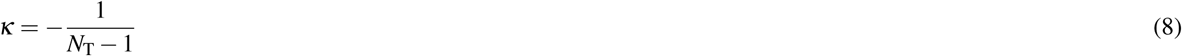

(ref.^34^, Eq. B.1). Contrastingly, in an island model with *n*_d_ demes with *N* individuals each (so that *N*_T_ = *n*_d_*N*) and a Moran demography (where adults have a positive probability of surviving to the next generation) scaled relatedness becomes

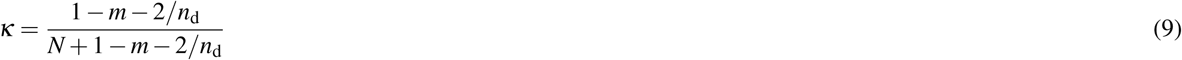

where *m* is the migration rate (ref.^34^, Eq. B.2). In this case, *κ* is inversely proportional to the migration rate *m*; it follows that an island model with less migration has, according to our definition, more spatial structure. As a final example, consider a transitive evolutionary graph of size *N*_T_ and degree *k* updated with a (death-birth) Moran demography. In this case we have

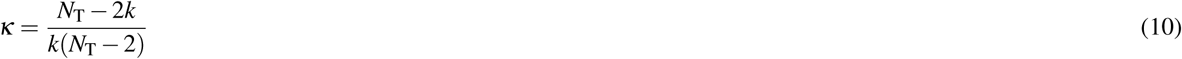

(ref.^34^, Eq. B.5 with *m* = 1; which can also be recovered from the value of *σ* given in Table 1 of ref.^43^ after applying the identity (7)). For this model of population structure, scaled relatedness is inversely proportional to the degree *k*. This means that, according to our terminology, graphs of larger degree (and hence more similar to a well-mixed population represented by a complete graph for which *k* = *N*_T_ − 1) are characterized by smaller spatial structure.

## Results

Calculating the gains from switching by first replacing Eq. (1) and (2) into Eq. (5) and (6), then replacing the resulting expressions into Eq. (4a) and (4b), and simplifying, we obtain that the gain function for the PGG with reward funds can be written as (see Methods)

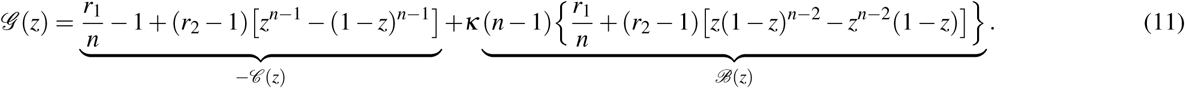

In the following, we identify convergence stable equilibria and characterize the evolutionary dynamics, first for well-mixed and then for spatially structured populations.

### Infinitely large well-mixed populations

For well-mixed populations, and as *N*_T_ → ∞, the scaled relatedness coefficient reduces to zero (Eq. (8)). In this case, the gain function simplifies to 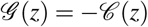 and we obtain the following characterization of the evolutionary dynamics (see Methods). If *r*_1_/*n* + *r*_2_ ≤ 2, *z* = 0 is the only stable equilibrium, and *RD* dominates *RC*. Otherwise, if *r*_1_/*n* + *r*_2_ *>* 2, both *z* = 0 and *z* = 1 are stable, and there is a unique *z*\* > 1/2 that is unstable. In this case, the evolutionary dynamics are characterized by bistability or positive frequency dependence, with the basin of attraction of full *RD* (*z* = 0) being always larger than the basin of attraction of full *RC* (*z* = 1). Moreover, *z*\* (and hence the basin of attraction of *z* = 0) decreases with increasing *r*_1_ and *r*_2_. In particular, higher reward funds lead to less stringent conditions for *RC* to evolve. In any case, *RC* has to be initially common (*z* > 1/2) in order for full *RC* to be the final evolutionary outcome.

### Spatially structured populations

Interactions in spatially structured populations (for which *κ* is not necessarily equal to zero) can dramatically alter the evolutionary dynamics of public goods with prosocial and antisocial rewards. In particular, we find that whether or not the extreme points *z* = 0 and *z* = 1 are stable depends on how the scaled relatedness coefficient *κ* compares to the critical values

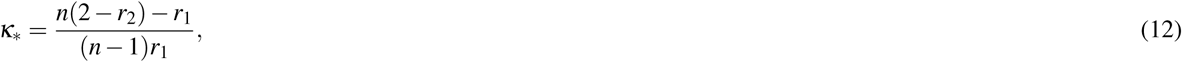

and

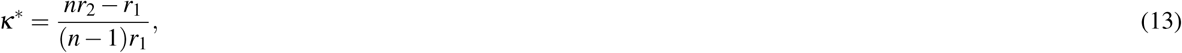

which satisfy *κ*_*_ ≤ *κ*\*, in the following way (Fig. 1):

1. For low values of *κ* (*κ* < *κ*_*_), full *RD* (*z* = 0) is stable and full *RC* (*z* = 1) is unstable.
2. For intermediate values of *κ* (*κ*_*_ < *κ* < *κ*\*), both full *RD* and full *RC* are stable.
3. For large values of *κ* (*κ* > *κ*\*), full *RC* is stable and full *RD* is unstable.

**Figure 1.**
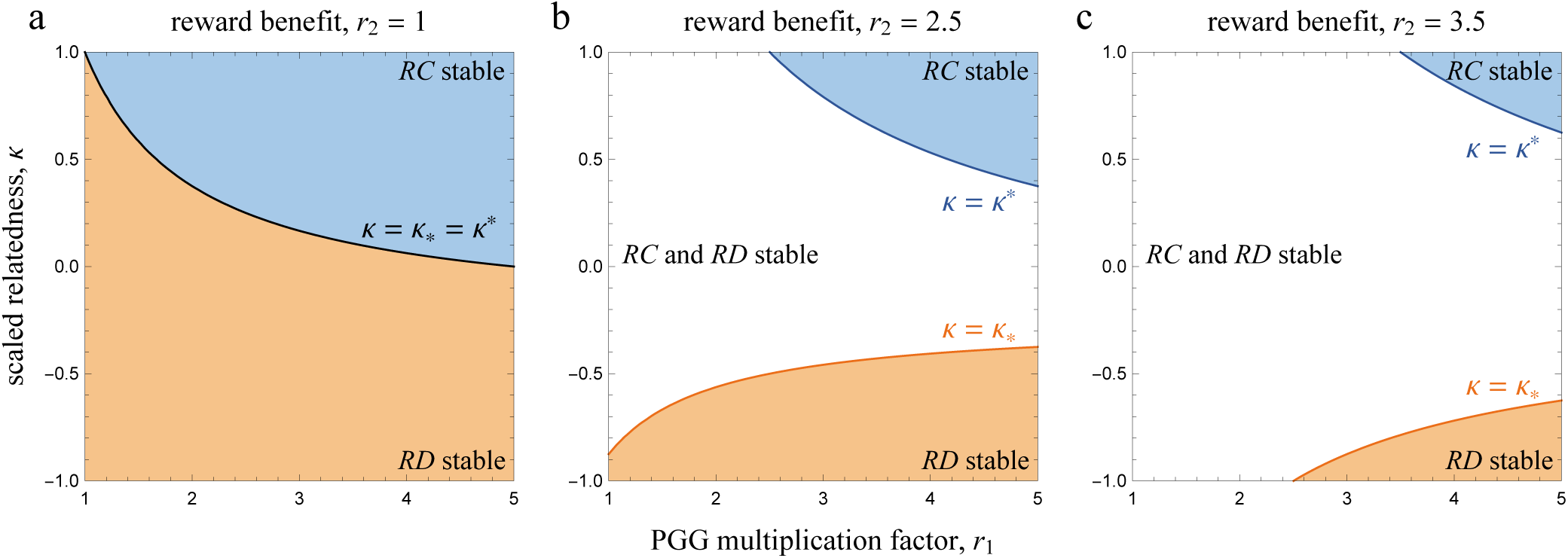
Phase diagrams illustrating the possible dynamical regimes of public goods games with prosocial and antisocial reward funds. Prosocial rewarding (*RC*) is stable if *κ > κ*_*_, while antisocial rewarding (*RD*) is stable if *κ < κ*\*. The critical values *κ*_*_ and *κ*\* are functions of the public goods game multiplication factor *r*_1_, the reward benefit *r*_2_, and the group size *n*, as given by Eqs. (12) and (13). Increasing the reward benefit *r*_2_ makes it more difficult for both prosocial and antisocial rewarding to increase from rare. Parameters: *n* = 5.

For a given group size *n* and PGG multiplication factor *r*_1_, *κ*_*_ = *κ*\* if and only if *r*_2_ = 1, i.e., if rewards are absent. In this case, full *RD* and full *RC* cannot be both stable.

Rewards have contrasting effects on *κ*_*_ (the critical scaled relatedness value below which full *RC* is unstable) and *κ*\* (the critical scaled relatedness value above which full *RD* is unstable). On the one hand, *κ*_*_ is decreasing in the reward benefit *r*_2_, so larger rewards increase the parameter space where full *RC* is stable. If spatial structure is maximal, i.e., *κ* = 1, the condition for full *RC* to be stable is *r*_1_ +*r*_2_ > 2, which always holds. On the other hand, *κ*\* is an increasing function of *r*_2_. Hence, larger rewards make it harder for spatial structure to destabilize the full *RD* equilibrium, and hence for *RC* to increase when rare. For *κ* = 1, full *RD* is still stable whenever *r*_1_ < *r*_2_. Contrastingly, full defection can never be stable if *κ* = 1 in the absence of rewards (i.e., *r*_2_ = 1) since, by definition, *r*_1_ > 1. From this analysis we can already conclude that even maximal spatial structure does not necessarily allow *RC* to invade and increase when rare. In addition, a minimum value of scaled relatedness is required for prosocial rewarding to be stable once it is fully adopted by the entire population.

Let us now investigate singular strategies. Depending on the parameter values, there can be either zero, one, or three interior points at which the gain function (and hence the selection gradient) vanishes (see Methods). If there is a unique singular point, then it is unstable while *z* = 0 and *z* = 1 are stable, and the evolutionary dynamics is characterized by bistability. If there are three singular points (probabilities *z*_L_, *z*_M_, and *z*_R_, satisfying 0 < *z*_L_ < *z*_M_ < *z*_R_ < 1), then *z* = 0, *z*_M_, and *z* = 1 are stable, while *z*_L_ and *z*_R_ are unstable. In this case *RD* and *RC* coexist at the convergence stable mixed strategy *z*_M_; a necessary condition for this dynamical outcome is both relatively large reward benefits and relatively large scaled relatedness.

We calculated the singular strategies numerically, as the equation 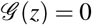 cannot be solved algebraically in the general case (Fig. 2 and Fig. 3). Increasing scaled relatedness generally increases the parameter space where *RC* is favored. Yet, there are cases where increasing scaled relatedness can hinder the evolution of *RC*. Specifically, when the reward benefit is considerably larger than the public goods share, increasing scaled relatedness can increase the basin of attraction of the full *RD* equilibrium (Fig. 2*c*). Also, increasing rewards can be detrimental to *RC* in spatially structured populations by increasing the basin of attraction of full *RD* (Fig. 3*c*, *f*, *h*, *i*); this is never the case when there is no spatial structure (Fig. 3*a*, *d*, *g*). Finally, the best case scenario from the point of view of a rare mutant playing *z* = *δ* (where *δ* is vanishingly small) is in the absence of rewards (i.e., *r*_2_ = 1), because that is the case where the required threshold value of scaled relatedness to favor prosocial rewarding is the lowest (i.e., where *κ*\* attains its minimum value in Eq. (13)).

**Figure 2.**
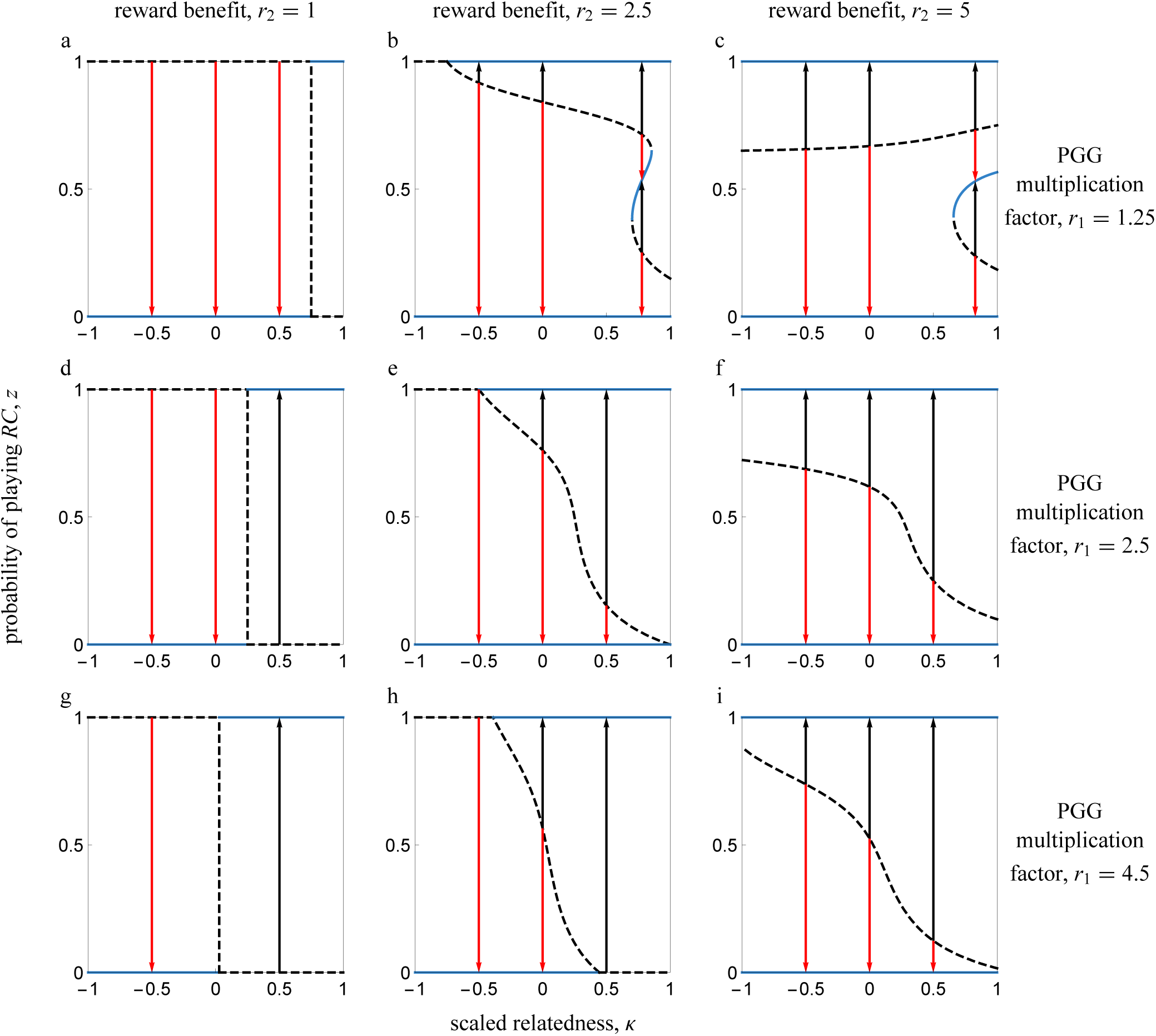
Bifurcation plots illustrating the evolutionary dynamics of pool rewarding in spatially structured populations. The scaled relatedness coefficient serves as a control parameter. Arrows show the direction of evolution for the probability of playing prosocial rewarding. Solid (dashed) lines correspond to convergence stable (unstable) equilibria. In the left column panels (*a*, *d*, *g*), rewards are absent (i.e., *r*_2_ = 1). In the middle column panels (*b*, *e*, *h*), *r*_2_ = 2.5. In the right column panels (*c*, *f*, *i*), *r*_2_ = 4.5. In the top row panels (*a*, *b*, *c*), *r*_1_ = 1.25. In the middle row panels (*d*, *e*, *f*), *r*_1_ = 2.5. In the bottom row panels (*g*, *h*, *i*), *r*_1_ = 4.5. In all panels, *n* = 5. A value of *κ* = 0 could correspond to an infinitely large well-mixed population (Eq. (8)); a value of *κ* = 0.25 could correspond to an evolutionary graph updated with a death-birth Moran model with *N*_T_ ≫ *k* and *k* = 4 (Eq. (10)); a value of *κ* ≈ 0.167 could correspond to an infinite island model with deme size *N* = 5 and *m* ≪ 1 (Eq. (9)).

**Figure 3.**
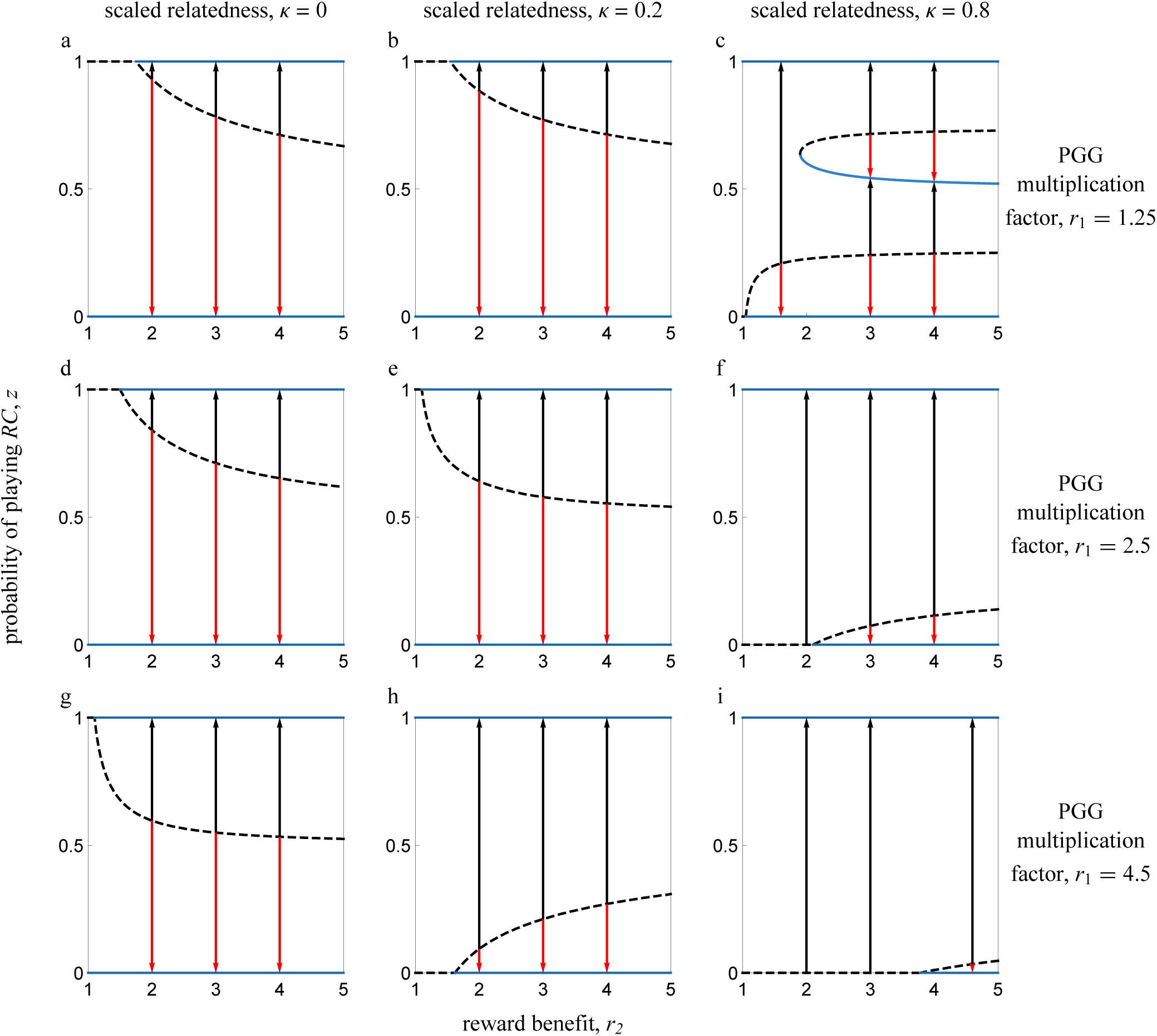
Bifurcation plots illustrating the evolutionary dynamics of pool rewarding in spatially structured populations. The reward benefit serves as a control parameter. Arrows show the direction of evolution for the probability of playing prosocial rewarding. Solid (dashed) lines correspond to convergence stable (unstable) equilibria. In the left column panels (*a*, *d*, *g*), there is no spatial structure (i.e., *κ* = 0). In the middle column panels (*b*, *e*, *h*), *κ* = 0.2. In the right column panels (*c*, *f*, *i*), *κ* = 0.8. In the top row panels (*a*, *b*, *c*), *r*_1_ = 1.25. In the middle row panels (*d*, *e*, *f*), *r*_1_ = 2.5. In the bottom row panels (*g*, *h*, *i*), *r*_1_ = 4.5. In all panels, *n* = 5.

In order to understand why, contrary to naive expectations, increasing spatial structure might sometimes select against *RC*, note first that the derivative of the gain function with respect to *κ* is equal to the indirect effect 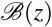. This is nonnegative if

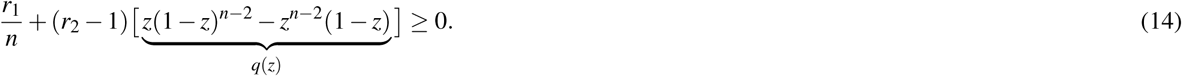

In the absence of rewards (i.e., *r*_2_ = 1), condition (14) always holds. That is, increasing scaled relatedness always promotes cooperation when there are no rewards. In addition, when 0 ≤ *z* ≤ 1/2, the function *q*(*z*) is nonnegative, so that condition (14) holds and 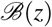 is positive. Hence, increasing scaled relatedness is always beneficial for *RC* when such behavior is expressed less often than *RD*. However, increasing scaled relatedness might not always favor *RC* when such behavior is already common in the population, i.e., if *z* > 1/2. Indeed, when the multiplication factor of the PGG is relatively small and rewards are relatively large, condition (14) is not fulfilled for some *z* and 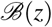 is negative for some probability of playing *RC* (Fig. 4).

**Figure 4.**
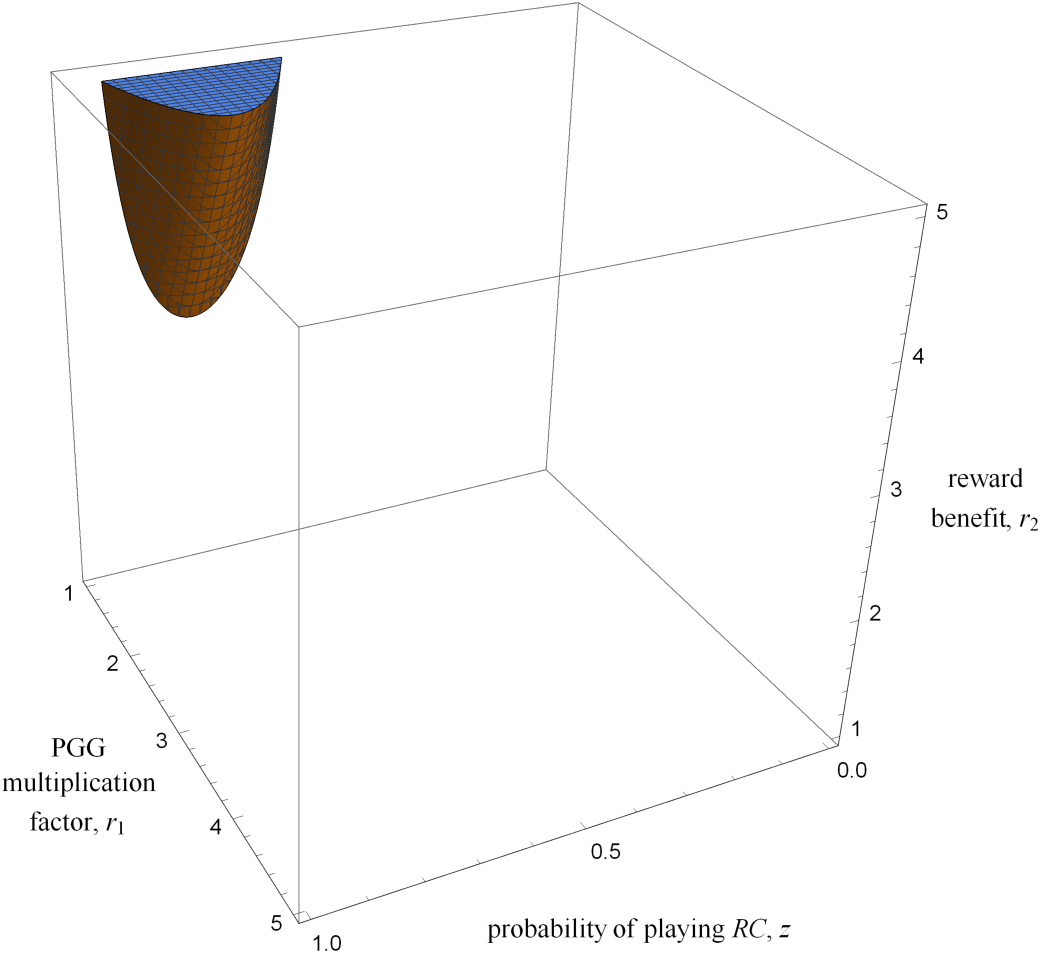
Parameter space where condition (14) does not hold and increasing spatial structure is detrimental to prosocial rewarding for some values of the probability of playing prosocial rewarding, *z*. Parameters: *n* = 5.

A closer look at the indirect gains from switching Θ*_k_* (Eq. 6) reveals why 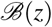, and hence the effect of scaled relatedness on the selection gradient, can be negative for some *z*. The indirect gains from switching are nonnegative for all *k* ≠ *n*−2. For *k* = *n*−2 and *n* ≥ 4 we have Θ*_n_*_−2_ = (*n*−1)*r*_1_/*n*−*r*_2_+1, which can be negative if

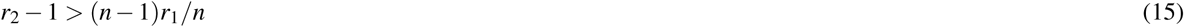

holds. Inequality (15) is hence a necessary condition for 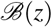 to be negative for some *z* and for prosocial rewarding to fail to qualify as payoff cooperative or payoff altruistic [*sensu* ref.^36^]. Indeed, when condition (15) holds and hence Θ*_n_*_−2_ *<* 0, prosocial rewarding cannot be said to be altruistic according to the “focal-complement” interpretation of altruism^57,58^. This is because the sum of the payoffs of the *n*−1 co-players of a given focal individual, out of which *n*−2 play *RC* and one plays *RD*, is larger if the focal plays *RD* than if the focal plays *RC*. We also point out that *RC* is not altruistic according to an “individual-centered” interpretation^58,59^ or “cooperative” [*sensu* ref.^60^] if

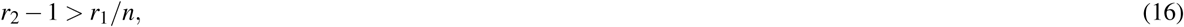

since in this case the payoff to a focal individual playing *RD* as a function of the number of other players choosing *RC* in the group, *d_k_* (see Eq. (2)), is decreasing (and not increasing) with *k* at *k* = *n*−2. Indeed, if condition (16) holds, players do not necessarily prefer other group members to play *RC* irrespective of their own strategy: a focal *RD* player would prefer one of its *n* − 1 co-players to play *RD* rather than play *RC*. In the light of this analysis, it is perhaps less surprising that for some parameters increasing spatial structure can be detrimental to the evolution of prosocial rewarding, even if prosocial rewarding Pareto dominates antisocial rewarding.

### Individual-based simulations

To test the validity of our mathematical model, we also ran individual-based simulations for a well-mixed population and a square lattice of size *N*_T_ = 400, both updated with a Moran death-birth rule (see Methods for details). As predicted by our mathematical analysis, the evolutionary dynamics are often characterized by a single interior convergence unstable point *z*\* (Fig. 5). When the phenotypic value *z*_0_ of the initially monomorphic population is below such point, selection tends to disfavors rewarding cooperation and the population converges to full *RD* (*z* = 0). Contrastingly, when the population starts with a phenotypic value larger than *z*\*, selection tends to favor rewarding cooperation and the population converges to full *RC* (*z* = 1). In all cases, the convergence unstable strategy *z*\* resulting from our mathematical analysis are very good predictors of the point at which selection changes direction in our individual-based simulations.

**Figure 5.**
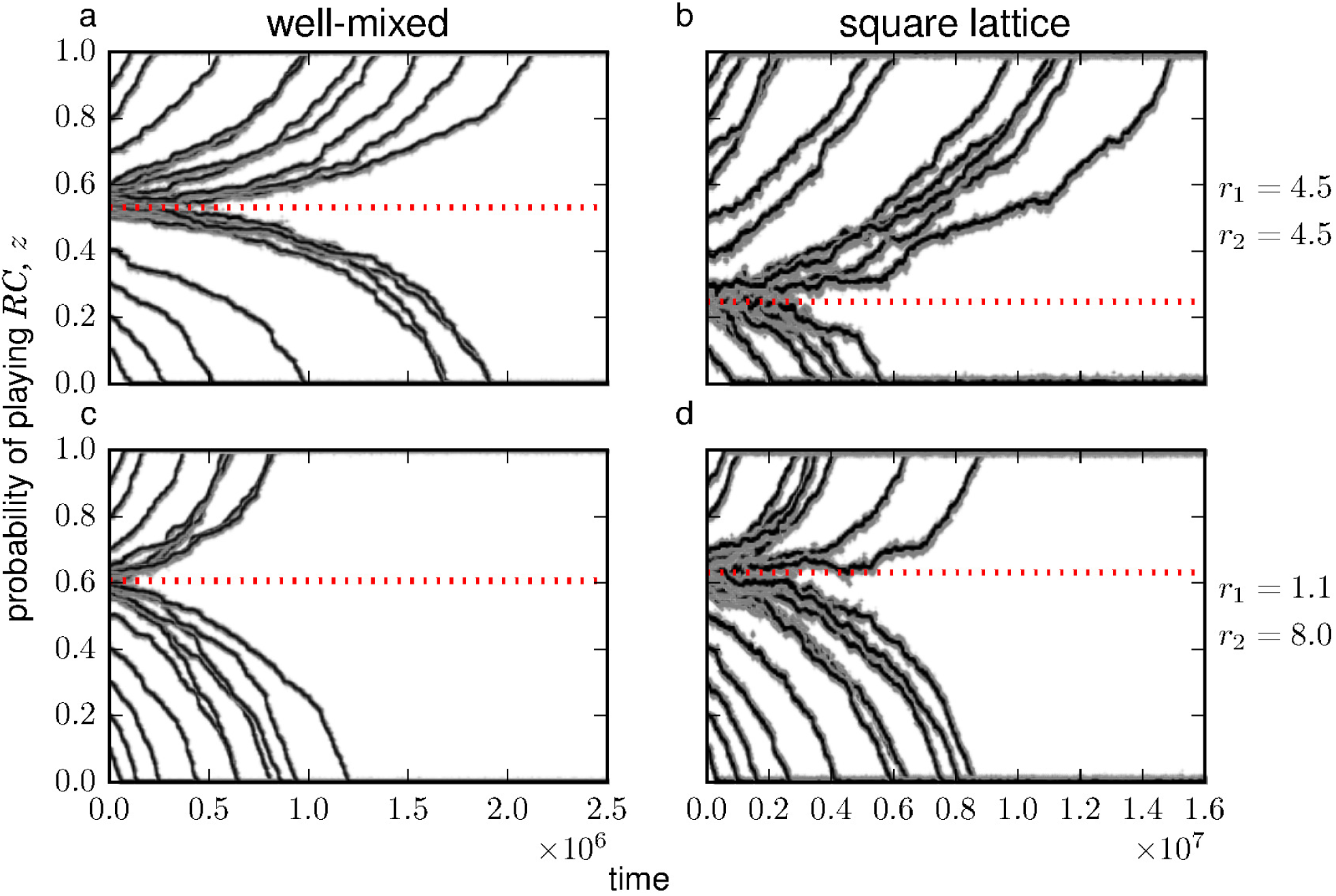
Evolution of the probability of playing rewarding cooperation in a simulated population of *N*_T_ = 400 individuals interacting in groups of size *n* = 5. Solid lines show the average phenotypic value of the population, gray dots show trait values of 10 individuals randomly sampled every *N*_T_ time steps. Each set of solid line and gray dots represent one realization of the stochastic process starting with a different initial condition where the population is monomorphic for trait value *z*_0_. Dotted red lines represent the analytical prediction for the value of the convergence unstable interior point *z*\*. Parameters: *w* = 10, *μ* = 0.01, *ν* = 0.05. Left panels (*a*, *c*): well-mixed population updated with a Moran death-birth process (*κ* ≈ −0.0025). Right panels (*b*, *d*): square lattice with periodic boundary conditions and von Neumann neighborhood, i.e., each node is connected to North, East, South, and West neighbors (*κ* ≈ 0.2462). Top row panels (*a*, *b*): *r*_1_ = 4.5, *r*_2_ = 4.5. Bottom row panels (*c*, *d*): *r*_1_ = 1.1, *r*_2_ = 8.0. In all cases, the analytical model predicts bistable evolutionary dynamics with a single convergence unstable equilibrium *z*\* dividing the basins of attraction of the two stable equilibria *z* = 0 and *z* = 1. a. *z*\* ≈ 0.5309. b. *z*\* ≈ 0.2472. c. *z*\* ≈ 0.606. d. *z*\* ≈ 0.632.

## Discussion

We have investigated the effect of spatial structure on the evolution of public goods cooperation with reward funds. Measuring spatial structure by means of a scaled relatedness coefficient allowed us to capture both the effects of increased genetic assortment and increased local competition that characterize evolution in spatially structured populations. We have found that (i) prosocial rewarding cannot invade full antisocial rewarding unless scaled relatedness is sufficiently large, but (ii) increasing scaled relatedness can be detrimental to prosocial rewarding in cases where rewards are considerably larger than the public goods share. We have also demonstrated the contrasting effects of increasing rewards, which (iii) only benefits prosocial rewarding in well-mixed populations, but (iv) can also benefit antisocial rewarding in spatially structured populations. These results illustrate how pool-rewards introduce non-linearities to the public goods game, which in turn lead to counterintuitive effects of population structure on the evolutionary dynamics.

We found that increasing spatial structure can sometimes be detrimental to prosocial rewarding. We confirmed this analytical result with individual-based simulations, where we showed that the basin of attraction of full prosocial rewarding can be greater in a well-mixed population than in a square lattice with a Moran death-birth demography. This is a somewhat counterintuitive result, because spatial structure often favors the evolution of cooperation^39,41,50,60–62^. However, previous studies have shown that increasing population structure can sometimes have a negative effect on the evolution of cooperative strategies^34,63,64^. For example, Hauert and Doebeli^63^ found that, when social interactions are modelled as a two-player snowdrift game between pure strategists under strong selection, well-mixed populations lead to higher levels of cooperation than square lattices. Peña *et al.*^64^ studied a multiplayer version of this game, but assuming weak selection and a death-birth Moran process, and found similar results. In our case, spatial structure can oppose prosocial rewarding because the indirect gains from switching from antisocial to prosocial rewarding can be negative. In particular, if individuals interact in groups of size *n* ≥ 4 and exactly *n* − 2 co-players choose rewarding cooperation while one co-player chooses rewarding defection, playing rewarding defection (*RD*) rather than rewarding cooperation (*RC*) might increase (rather than decrease) the payoffs of co-players. The reason is that, in this case, by choosing *RD* the focal player helps its *RD* co-player getting the reward fund, while also allowing its *RC* co-players to keep theirs, as the focal contribution is not critical to the creation of the prosocial reward fund. If the reward benefit is so large that Eq. (15) holds, the benefit to the single *RD* co-player is greater than what everybody loses by the focal not contributing to the public good, and the sum of payoffs to co-players is greater if the focal plays *RD* than if it plays *RC*. This implies that, although prosocial rewarding Pareto dominates antisocial rewarding, it does not strictly qualify as being payoff altruistic or payoff cooperative^36^, hence the mixed effects of increasing spatial structure.

Our results also revealed the fact that higher values of the reward benefit *r*_2_ make it more difficult for prosocial rewarding to invade from rare in spatially structured populations. Indeed, the critical value of scaled relatedness required for prosocial rewarding to be favored over antisocial rewarding is greater in the presence of rewards than in their absence. This is important because rewards are meant to be mechanisms incentivizing provision in public goods games^15,18^, rather than making collective action more difficult to emerge. Clearly, our result hinges on the assumption that prosocial rewarders and antisocial rewarders are both equally effective in rewarding themselves, i.e., that *r*_2_ is the same for both prosocial and antisocial rewarders. Challenging this assumption by making investments in rewards contingent on the production of the public good, or by increasing the ability of prosocials to reward each other relative to that of antisocials^19^, will necessarily change this picture and promote prosocial rewarding in larger regions of the parameter space.

Although higher rewards prevent the invasion of prosocial rewarding from rare, we have also shown that, once prosocial rewarding is common, higher rewards can further enhance the evolution of prosocial rewarding. These results are in line with the findings of SP15, who showed that when both spatial structure is sufficiently large (their spatial model supports cooperation even in the absence of rewards) and the initial frequency of prosocial rewarding is relatively high (i.e., 1/4 in their simulations), larger rewards promote prosocial rewarding.

In contrast to the original model by SP15, which considered discrete strategies and strong selection, we assumed continuous mixed strategies and weak selection^36^. For well-mixed populations, it is well known that these different sets of assumptions lead to identical results under a suitable reinterpretation of the model variables^53^. Thus, our result that in this case there is a unique convergence unstable *z*\* in mixed strategies also implies that the replicator dynamics for the two-strategy model will be characterized by an unstable rest point at a frequency *z*\* of prosocial rewarders, and corroborates the numerical results presented in section 3.a of SP15. By contrast, for structured populations the invasion and equilibrium conditions between discrete- and mixed-strategy game models^45,46,63^ and between weak and strong selection models^61^ can differ. Hence, our results for spatially structured populations need not be identical to those reported in SP15. Importantly, our mathematical framework assumes that the population is essentially monomorphic. An initial state of, say, *z* = 1/3 in our Fig. 2 means that all individuals in the population play the same mixed strategy *z* = 1/3. Evolution then proceeds by means of a trait substitution sequence (TSS), whereby a single mutant (which we also assume plays a slightly different mixed strategy *z* + *δ*, where *δ* is small) will either become extinct or invade and replace the resident population^65^. If the latter happens, the resident strategy is updated to *z* + *δ* and the process starts again, until a convergence stable state is reached. The TSS assumption, common in adaptive dynamics and related mathematical methods studying the evolution of continuous traits in spatially structured populations^28,66^ is then in stark contrast to the numerical simulations used in SP15 and related studies^41^, where evolution starts from a polymorphic population where a large number of mutants appear en masse either randomly or clustered together according to a given “prepared initial state”.

Finally, there is a slight difference in the way SP15 and our study (explicitly in our simulations and implicitly in our analytical model) implemented the evolutionary game on a graph. SP15 assume, in the tradition of other computational studies (e.g., refs^41,67,68^), that a focal player’s total payoff is given by the sum of payoffs obtained in *k* + 1 different games, one “centered” on the focal player itself and the other *k* centered on its neighbors. As a result, a focal player interacts not only with first-order but also with second-order neighbors. An analytical treatment of such case would need an extension of our framework to incorporate an additional scaled relatedness coefficient to account for interactions with second-order neighbors^50^. Contrastingly, we assumed that a focal player obtains its payoff from a single multiplayer game with its *k* immediate neighbors^64,69^. This more parsimonious assumption allows us to analyze multiplayer interactions on graphs in a straightforward way (i.e., by allowing us to directly use the framework developed in ref.^36^ requiring a single scaled relatedness coefficient) without importantly modifying the underlying evolutionary dynamics^69^.

Our motivation for a set of assumptions different to those of SP15 was hence for both analytical tractability and wider applicability. An analytical solution of the model with discrete strategies (as in SP15) would require tracking higher-order genetic associations and effects of local competition, which can be a complicated task even in relatively simple models of spatial structure under weak selection^61,70–73^. By contrast, assuming continuous strategies (and a single game per player) allowed us to identify convergence stable levels of prosocial rewarding in a wide array of spatially structured populations, each characterized by a particular value of scaled relatedness. This way, we made analytical progress going beyond the numerical results on a particular type of population structure (a square lattice with overlapping groups of size *n* = 5) studied in SP15. Furthermore, treating scaled relatedness as an exogenous parameter independent of specific demographic assumptions, allows making more general predictions on the evolution of a trait, as well as relating results from different types of models^74^. Understanding the effect of more specific demographic parameters (such as the dispersal rate, the update rule, or the degree of the network) can be achieved by determining how scaled relatedness can be expressed in terms of those parameters, as we have shown above.

Our analytical results are valid only to the first order of *δ* (the difference between the trait of mutants and the trait of residents). As a result, we cannot evaluate whether or not the singular strategies we identify as convergence stable are also “evolutionarily stable” or “locally uninvadable”, i.e., whether a population monomorphic for a singular value will resist invasion by mutants with traits close to the singular value^37,52,54,75^. This also means that our model does not allow one to check whether or not evolutionary branching (whereby a convergence stable but locally invadable population diversifies into differentiated coexisting morphs^76^) might occur. We hasten to note that such drawback is not particular to our method^28^. Evolutionary stability in spatially structured populations is significantly more challenging to characterize than convergence stability^37,77–80^ and is thus beyond the scope of the present paper.

A related issue has to do with our assumption that individuals play mixed strategies and hence that payoffs are linear in the focal’s own strategy. For this kind of models, “a peculiar degeneracy raises its ugly head”^81^, namely that the second-order condition to evaluate evolutionary stability in a well-mixed population is null. In turn, this implies that phenotypic variants at a singular point that is convergence stable are strictly neutral. Such degeneracy is however restricted to well-mixed populations, and does not necessarily apply to spatially structured populations. Indeed, the condition for uninvadibility under weak selection in subdivided populations has been shown to depend also on mixed partial derivatives of the payoff function^37^, which in general are not zero. All in all, our view is that assuming individuals play mixed strategies of a matrix game is not that problematic: For well-mixed populations (where the degeneracy raises its head), the convergence stable mixed strategies can be reinterpreted as evolutionarily stable points of a replicator dynamics in discrete strategies; for spatially structured populations, there is simply no degeneracy. Future work should explore the conditions under which convergence stable mixed strategies of the model presented here and other matrix games are locally uninvadable.

Both our model and that of SP15 have not considered the presence of individuals who are able to benefit from reward funds without contributing to them. In other words, second-order defection is avoided by design. Allowing for second-order defection makes cooperation through pool rewarding vulnerable, even in the absence of antisocial rewarding^15^. Therefore, even though the conclusions of SP15 contradict the findings of dos Santos^19^, namely that antisocial rewarding deters cooperation except in certain conditions (e.g., better rewarding abilities for prosocials), SP15 did not investigate standard pool-rewarding^15,18,19^. Hence, their claim that spatial structure prevents antisocial rewarding from deterring cooperation, while not always true as we have shown here, does not apply to the more general case of pool-reward funds where second-order defection is allowed. Exploring the effects of spatial structure in these more realistic cases remains an interesting line of research.

To conclude, we find that antisocial rewarding deters the evolution of cooperation from rare unless scaled relatedness is sufficiently high and rewards are relatively low, or ideally absent. We argue that additional countermeasures, such as exclusion and better rewarding abilities for prosocials^19^, are still required to (i) prevent antisocial rewarding from deterring cooperation between unrelated social partners, and (ii) allow prosocial rewarding to invade either when relatedness is low or when rewards are too large.

## Methods

### Gain function

To derive the gain function 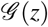 (Eq. (11)), we first calculate the direct and indirect gains from switching (Eqs. (5) and (6)) associated to the payoffs of the game. We find that the gains from switching depend on the group size *n* in the following way.

1. For *n* = 2: (Δ_0_, Δ_1_) = (*r*_1_/2 − *r*_2_*, r*_1_/2 + *r*_2_ − 2) and (Θ_0_, Θ_1_) = (*r*_1_/2 − *r*_2_ + 1*, r*_1_/2 + *r*_2_ − 1).
2. For *n* = 3: (Δ_0_, Δ_1_, Δ_2_) = (*r*_1_/3 − *r*_2_*, r*_1_/3 − 1*, r*_1_/3 + *r*_2_ − 2) and (Θ_0_, Θ_1_, Θ_2_) = (2*r*_1_/3, 2*r*_1_/3, 2*r*_1_/3).
3. For *n* = 4: (Δ_0_, Δ_1_, Δ_2_, Δ_3_) = (*r*_1_/4 − *r*_2_*, r*_1_/4 − 1, *r*_1_/4 − 1, *r*_1_/4 + *r*_2_ − 2) and (Θ_0_, Θ_1_, Θ_2_, Θ_3_) = (3*r*_1_/4, 3*r*_1_/4 + *r*_2_ − 1, 3*r*_1_/4 − *r*_2_ + 1, 3*r*_1_/4).
4. For *n* ≥ 5: (Δ_0_, Δ_1_,…, Δ*_n_*_−2_, Δ*_n_*_−1_) = (*r*_1_/*n*−*r*_2_*, r*_1_/*n*−1,…, *r*_1_*/n*−1*, r*_1_/*n*+*r*_2_ −2) and (Θ_0_, Θ_1_, Θ_2_,…, Θ*_n_*_−3_, Θ*_n_*_−2_, Θ*_n_*_−1_) = ((*n* − 1)*r*_1_/*n*, (*n* − 1)*r*_1_/*n* + *r*_2_ − 1, (*n* − 1)*r*_1_/*n*,…, (*n* − 1)*r*_1_/*n*, (*n* − 1)*r*_1_/*n* − *r*_2_ + 1, (*n* − 1)*r*_1_/*n*).

Replacing the direct gains from switching Δ*_k_* into the expression for the direct effect 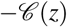 (Eq. (4a)) and the indirect gains from switching Θ*_k_* into the expression for the indirect effect 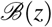 (Eq. (4b)), and simplifying, we obtain the gain function 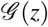 given in Eq. (11), which is valid for all *n* ≥ 2.

### Evolutionary dynamics for *κ* = 0

For *κ* = 0, the gain function 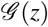 (Eq. (11)) reduces to 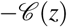 (Eq. (4a)). This function is increasing and its end-points are given by 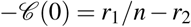 and 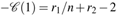. Since 1 < *r*_1_ < *n* and *r*_2_ > 1, 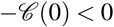 always holds and *z* = 0 is always stable. If *r*_1_*/n*+*r*_2_ ≥ 2, 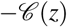 is nonpositive for all *z* and *z* = 0 is the only stable equilibrium. If *r*_1_*/n*+*r*_2_ < 2, 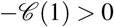 and *z* = 1 is also stable. In this case, and since 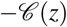 is increasing, 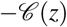 has a single zero *z*\* in (0, 1) corresponding to an unstable equilibrium. Such zero is given by the unique solution to

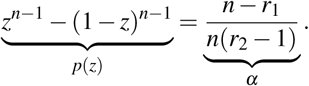

Since *p*(*z*) is increasing in *z*, *p*(1/2) = 0, and *α >* 0 always holds, it follows that *z*\* *>* 1/2. Additionally, since *α* is decreasing in both *r*_1_ and *r*_2_, *z*\* is increasing in both *r*_1_ and *r*_2_.

### Evolutionary dynamics for *κ* ≠ 0

Rearranging terms, the gain function 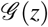 given by Eq. (11) can be alternatively written as

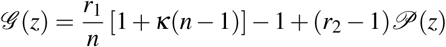

where

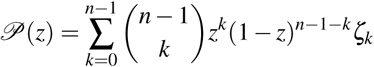

is a polynomial in Bernstein form^53^ of degree *n* − 1 with coefficients given by

1. (*ζ*_0_*, ζ*_1_) = (−(1 + *κ*), 1 + *κ*) if *n* = 2.
2. (*ζ*_0_*, ζ*_1_*, ζ*_2_) = (−1, 0, 1) if *n* = 3.
3. (*ζ*_0_*, ζ*_1_*, ζ*_2_*, ζ*_3_) = (−1*, κ,* −*κ*, 1) if *n* = 4.
4. (*ζ*_0_*, ζ*_1_*, ζ*_2_,…, *ζ_n_*_−3_, *ζ_n_*_−2_, *ζ_n_*_−1_) = (−1*, κ*, 0,…, 0, −*κ*, 1) if *n* ≥ 5.

The number of sign changes (and hence of singular points) of 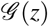 is bounded from above by the number of sign changes of 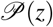. Moreover, and by the variation-diminishing property of polynomials in Bernstein form^53^, the number of sign changes of 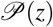 is equal to the number of sign changes of the sequence of coefficients (*ζ*_0_,…,ζ*_n_*_−1_) minus an even integer. It then follows that the number of singular points is at most one if *n* ≤ 3 or if scaled relatedness is nonpositive, *κ* ≤ 0. In this case, the unique singular point *z*\* is convergence unstable. However, if *n* ≥ 4 and *κ* > 0, there could be up to three singular points *z*_L_, *z*_M_, and *z*_R_ satisfying 0 < *z*_L_ < *z*_M_ < *z*_R_ < 1 such that *z*_L_ and *z*_R_ are convergence unstable and *z*_M_ is convergence stable.

### Computer simulations

We performed individual-based simulations for a population composed of *N*_T_ individuals, using NumPy version 1.11.1. Starting with a population monomorphic for a probability *z* = *z*_0_ of playing rewarding cooperation, we track the evolution of the phenotypic distribution as mutations of small effect continuously arise. Each individual *j* = 1*,…,N*_T_ is characterized by its probability *z_j_*. The average payoff of individual *j* is given by 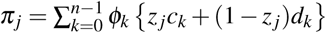 where the payoff sequences *c_k_* and *d_k_* are respectively given by Eqs. (1) and (2), and *ϕ_k_* is the probability that exactly *k* out of *n* − 1 of *j*’s neighbors choose action *RC*. Technically, *ϕ_k_* is the probability mass function of a Poisson binomial distribution with parameters 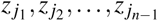, where *j*_ℓ_ represents the ℓ-th neighbor of individual *j*, and 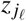 its phenotype. For simplicity, we approximate *ϕ_k_* by the probability mass function of a binomial random variable with parameters *n* and 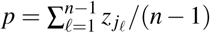; such approximationtothetrue probability mass function is very accurate when the probabilities 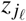 are close to each other. We update the population via a death-birth Moran process, i.e., we (i) randomly select one individual to die and one of its neighbors to give birth, and (ii) fill the vacated breeding spot with probability proportional to the average payoff. More specifically, assuming that *j* is chosen to die, its ℓ-th neighbor is given a probability 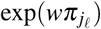 of replacing *j*, where *w* is the intensity of selection and 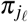 is the payoff of the ℓ-th neighbor of *j*. The vacated breeding spot is filled with the same phenotypic value as the parent with probability 1 − *μ* and with a mutated phenotypic value with probability *μ*. If mutation occurs, the mutated phenotypic value is given by the parent value plus a small perturbation sampled from a normal distribution with mean zero and standard deviation *ν*. The resulting phenotypes are truncated so that they are numbers between 0 and 1. The previous procedure is repeated for a sufficiently large number of time steps.

We simulated two models of spatial population structure: a square lattice with periodic boundary conditions (i.e., a type of transitive graph with *k* = 4) of *N*_T_ = 400 nodes, and a well-mixed population of *N*_T_ = 400 individuals where a given focal individual is assigned four randomly chosen other individuals as neighbors each time step. Other parameter values used were: *w* = 10, *μ* = 0.01, *ν* = 0.005.

## Acknowledgements

We thank Pat Barclay, Laurent Lehmann, and Stu West for comments, and the Swiss National Science Foundation for funding(Grant P2LAP3-158669 to M.D.S.).

## Author contributions statement

Both authors contributed equally to this work. M.D.S. and J.P. conceived and designed the study, performed the analysis, and wrote the manuscript. Both authors gave final approval for submission.

## Additional information

The authors declare no competing interests.

